# Representations of complex contexts: A role for hippocampus

**DOI:** 10.1101/766311

**Authors:** Halle R. Dimsdale-Zucker, Maria E. Montchal, Zachariah M. Reagh, Shao-Fang Wang, Laura A. Libby, Charan Ranganath

## Abstract

The hippocampus plays a critical role in supporting episodic memory, in large part by binding together experiences and items with surrounding contextual information. At present, however, little is known about the roles of different hippocampal subfields in supporting this item-context binding. To address this question, we constructed a task in which items were affiliated with differing types of context – cognitive associations that vary at the local, item level and membership in temporally organized lists that linked items together at a global level. Participants made item recognition judgments while undergoing high-resolution fMRI imaging. We performed voxel pattern similarity analyses to answer the question of how human hippocampal subfields represent retrieved information about cognitive states and the time at which a past event took place. As participants recollected previously presented items, activity patterns in the CA23DG subregion carried information about prior cognitive states associated with these items. We found no evidence to suggest reinstatement of information about temporal context at the level of list membership, but exploratory analyses revealed representations of temporal context at a coarse level in conjunction with representations of cognitive contexts. Results are consistent with characterizations of CA23DG as a critical site for binding together items and contexts in the service of memory retrieval.

## Introduction

Converging evidence suggests that the hippocampus (HC) plays a critical role in memory for events and their episodic details (Eichenbaum et al., 2007; Scoville & Milner, 1957; Vargha–Khadem et al., 1997). The HC is a circuit of interconnected subfields with different anatomical (Amaral & Witter, 1989; Burwell, 2000; Kesner & Rolls, 2015; Lavenex & Amaral, 2000; Witter et al., 2000), and, hence, computational (Marr, 1971; Norman & O’Reilly, 2003; Schapiro et al., 2017; Treves & Rolls, 1994) properties. Evidence from animal models has suggested that different hippocampal subfields uniquely contribute to memory through different computational specializations (e.g., pattern separation and completion; for reviews, see (Liu et al., 2015; Yassa & Stark, 2011) that enable one to encode the spatial and task variables that create a context for episodic memories (Kesner & Rolls, 2015; Levy, 1996; Lisman, 1999; Mankin et al., 2015; Norman & O’Reilly, 2003; Wallenstein et al., 1998). This is particularly true when task demands highlight the need to bind together item-in-context representations, such as learning where to search for a reward given an animal’s current location (McKenzie et al., 2014).

Evidence in humans generally corroborates the research in animal models, suggesting that, as a whole, the HC supports episodic memory by binding together information about items and the context in which they were encountered (Davachi, 2006; Eacott & Gaffan, 2005; Eichenbaum et al., 2007; O’Keefe & Nadel, 1978; Ranganath, 2010; Y. Zheng et al., 2021). At present, however, there is little known about the roles of different hippocampal subfields in item-context binding. Many studies have used high-resolution fMRI to examine item recognition (Bakker et al., 2008; Carr et al., 2010; Chen et al., 2011; Lacy et al., 2011; LaRocque et al., 2013; Reagh et al., 2014; Suthana et al., 2015; Viskontas et al., 2009; Yassa et al., 2011; Zeineh et al., 2003) or representations of spatial contexts (Brown et al., 2014, 2014; Chanales et al., 2017; Copara et al., 2014, 2014; Dimsdale-Zucker et al., 2018; Stokes et al., 2015; Suthana et al., 2011; L. Zheng et al., 2021).

Cognitive theories of episodic memory have conceptualized context in at least two ways. One view emphasizes context as a variable that changes over time (Estes, 1955; Howard & Kahana, 2002) due to ongoing fluctuations in the participant’s environment and cognitive state (e.g., the frank passage of time, changes in stimuli, shifting mental and physical states, etc.). Some high-resolution imaging studies have revealed evidence suggesting that the hippocampus can represent information about temporal context associated with particular items or associations between items based on temporal contiguity (Copara et al., 2014; Deuker et al., 2016; Dimsdale-Zucker et al., 2018; Nielson et al., 2015; Schapiro et al., 2013).

Another view is that episodic memories are associated with “cognitive contexts” (Diana et al., 2012, 2013)—relatively discrete cognitive states that are tied to attentional priorities, task demands, or currently relevant goals (Aly & Turk-Browne, 2016a, 2016b; Antony et al., 2021; Diana et al., 2008, 2012). These views are not mutually exclusive, and some models suggest that context in episodic memory may be represented through a combination of temporal context and cognitive representations of task states (Lohnas et al., 2015; Polyn et al., 2009).

In order to understand whether shifting cognitive contexts affiliated with encoding items could drive differences in representations across hippocampal subfields, we constructed a task in which we experimentally manipulated the cognitive context affiliated with studied items by varying the encoding question associated with an item (Davachi et al., 2003; Diana et al., 2008, 2012, 2013; Dzulkifli & Wilding, 2005; J. D. Johnson et al., 2009; M. K. Johnson et al., 1997; McDuff et al., 2009; Nolde et al., 1998; Polyn et al., 2009; Ranganath et al., 2004; Ritchey et al., 2015). To manipulate temporal context, we organized items into repeated lists, such that items in the same list would be expected to have stronger intra-list temporal associations than items in different lists. This allowed us to test the extent to which hippocampal subfields represent information about cognitive (i.e., encoding question) and temporal (i.e., list membership) contextual associations. We then scanned participants while they recollected these objects and used voxel pattern similarity (PS) analyses (Dimsdale-Zucker & Ranganath, 2018; Kriegeskorte et al., 2008) to examine patterns of activity in hippocampal subfields reflected spontaneous retrieval of information from the study phase about the temporal and cognitive context associated with each item.

## Method

### Participants

32 participants were recruited from the community and were compensated $50 for their time. This study was approved by the Institutional Review Board at the University of California, Davis. Four participants were excluded due to missing behavioral data, two participants were excluded for excessive motion that prevented tracing of hippocampal subfields, one participant was excluded due to an experimenter error at data collection that resulted in the incorrect stimuli being seen, and one participant was excluded because they only had one run of usable data after discarding motion-contaminated and data-collection contaminated runs. The results below reflect data from 24 remaining participants (Mage = 22.85 years, SD = 3.06 years, Nfemale = 13). One of these 24 participants was excluded from behavioral cognitive and temporal context analyses due to partially missing data; since the brain imaging data for this participant were complete and did not depend on this behavior being recorded, they were included in all other analyses.

### Encoding

Participants viewed eight 36-item lists of still pictures of everyday objects (e.g., contact lens case, french fries; http://cvcl.mit.edu/mm/uniqueObjects.html; see Figure 1). Object assignment to list and presentation order of objects within a list were uniquely randomized for each participant via the Matlab randperm function. To encourage participants to learn temporal relationships amongst items in a list (Palombo et al., 2019), each list was presented three times in a mini-block before subjects saw items from the next list (e.g., 1, 1, 1, 2, 2, 2…8, 8, 8). A label with a list number (e.g., “List 1”) appeared at the beginning of each list prior to seeing any objects. Mini-blocks were separated with a self-paced break. Presentation order of objects within a list was identical for all three list presentations.

**Figure 1.**
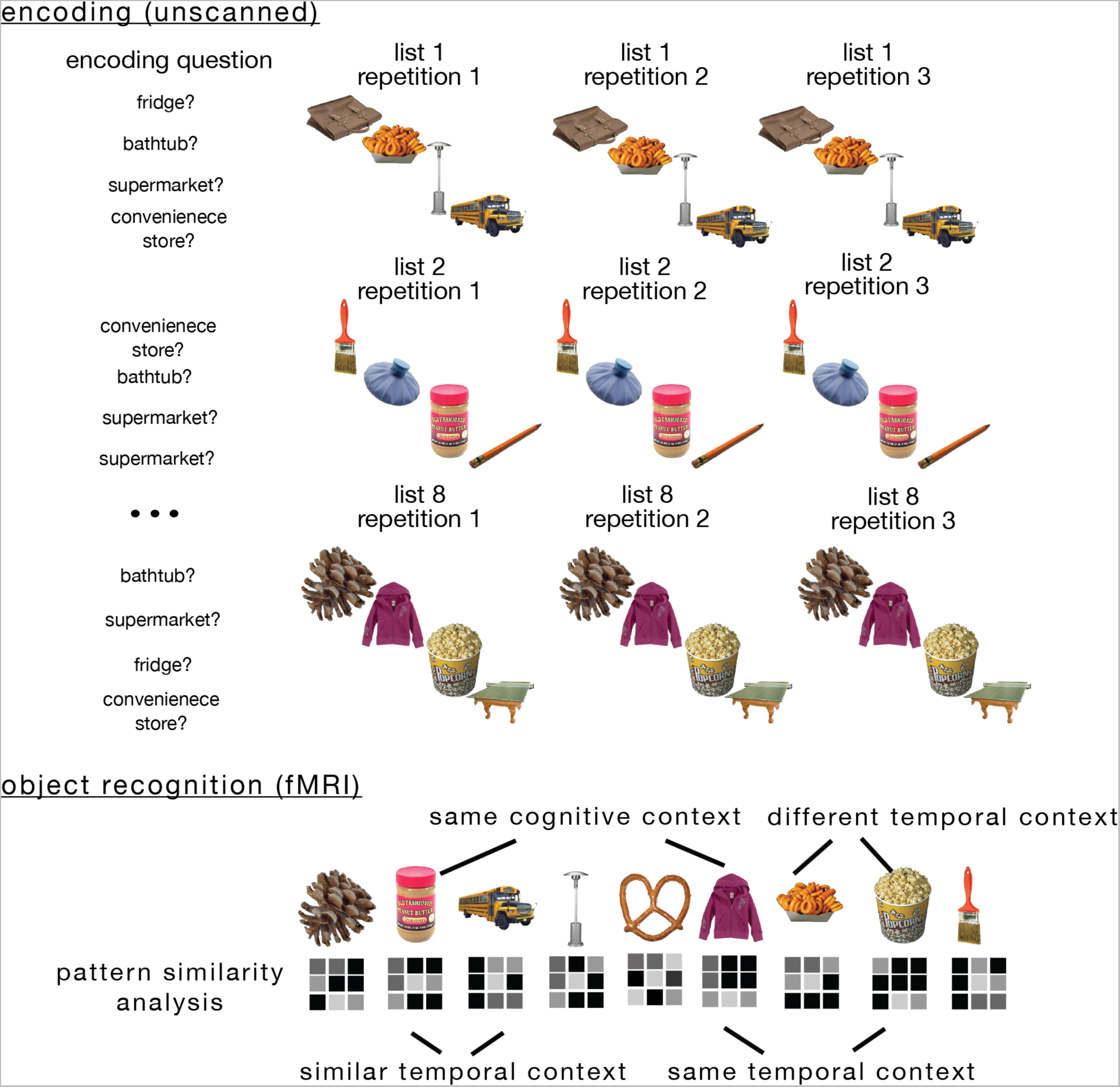
Task structure. During the encoding phase, each participant studied lists of 36 objects that were each randomly paired with one of four encoding questions (“Would this item fit in a fridge?”, “Would this item fit in a bathtub?”, “Would you find this item in a supermarket?”, “Would you find this item in a convenience store?”). Each list was repeated three times in a row to promote learning of the temporal relationships amongst the items. Objects appeared in the same order and with the same question (cognitive context) across all repetitions. High-resolution functional magnetic resonance brain imaging (fMRI) was used to examine hippocampal activity patterns during a recognition memory test for these objects, allowing us to examine activity pattern similarity as a function of whether pairs of items were encoded within the same or similar temporal contexts (i.e., studied in the same list or temporally proximal lists), and/or the same cognitive context (i.e., associated encoding question).

Objects remained on the screen for 2.5 seconds (timing and presentation parameters were controlled via Presentation [Neurobehavioral Systems, Inc., Berkeley, CA, www.neurobs.com]) while the participant made a yes/no button response to an orienting question (cognitive context). To manipulate cognitive context, each object was associated with one of four questions: Would this item fit in a refrigerator?, Would this item fit in a bathtub?, Would you find this item in a convenience store?, Would you find this item in a supermarket?. Each of the four questions was presented equally often in each block, and question/object pairs remained the same across all three list presentations. Participants were instructed that this was a decision-making task and that there would be some repetition but to concentrate on doing the task. Participants were not aware that memory for these questions would be tested later, thus the learning of question (cognitive) and temporal context information was incidental. Grouping items into lists was meant to induce a shared context representation for items in the same list. We reasoned that repetition of lists during the study phase would enhance contextual relationships between items in the same list. A similar temporal grouping strategy has been used previously to encourage connections between items in this fashion (Palombo et al., 2019). In contrast to the inter-item links facilitated by list membership, previous work has shown that interspersing questions throughout the encoding phase can result in fast, shifting variations in context that are aligned to changes in cognitive state (Polyn et al., 2009). Thus, to target cognitive context, each item was affiliated with one of four orienting questions (for a similar manipulation of cognitive context see (Diana et al., 2012). We note that because we did not collect fMRI data during the encoding phase, we are limited to looking at reactivated representations of cognitive and temporal contexts at the time of retrieval.

### Scanned object recognition

While in the MRI scanner, participants saw each of the 288 old objects from encoding as well as 72 new objects presented one at a time for 2.5 seconds with a jittered ITI ranging from 2-15 seconds (mean ITI jitter = 6 seconds). Objects were divided into 6 runs (60 trials per run). Object order within a run was pseudo-randomized such that objects with the same encoding question always had at least one intervening object (e.g., fridge, convenience store, bathtub, fridge, supermarket, fridge, etc.) to help minimize encoding context reinstatement biases on PS results (see Multivariate Results below). Proximity of objects from encoding mini-blocks (1-8) was not considered in the pseudo-randomization.

While in the scanner, participants were instructed to indicate via button press whether or not they remembered the object on a 4-point scale: 1=new, 2=familiar (old but no remembered details), 3=remembered non-temporal details (e.g., the encoding question, something about the object itself, or an association they made with the object), 4=remembered temporal detail (e.g., in what list or when they had seen the object during encoding). Responses for remembered judgments were collapsed into a single response bin for behavioral and fMRI analyses.

### Source memory: Cognitive context

After completing MRI scanning, participants returned to the lab where they completed a cognitive context source memory task. In this phase, participants saw all 288 studied objects from encoding and were asked to indicate which encoding question (fridge/bathtub/convenience store/grocery store) had been associated with the object. Objects were presented across four blocks of 72 trials each. Within each block, there were an equal number of objects from each encoding mini-block (1-8). Presentation order of objects was uniquely randomized by participant within each source memory block. Objects appeared on the screen until the participant had made their source memory judgment. There was no opportunity to guess or skip objects.

### Source memory: Temporal context

After completing the cognitive context source memory test, participants again saw the 288 old objects from encoding and this time were asked to indicate in which mini-block (1-8) the object had appeared. Objects were presented centrally on the screen and a number line displaying the numeric options 1-8 appeared below. Participants were told to use the numbers at the top of the keypad to make their response. Objects were again divided across four blocks of 72 trials with a different randomization order than was used in the task context source memory test. Objects remained on the screen until the participant had made their response. There was no opportunity to guess or skip temporal context judgments.

### fMRI acquisition and pre-processing

Scans were acquired on a Siemens Skyra 3T scanner with a 32 channel head coil. Two sets of structural images were acquired to enable subfield segmentation: A T1-weighted magnetization prepared rapid acquisition gradient echo (MP-RAGE) pulse sequence image (1 mm isotropic voxels), and a high-resolution T2-weighted image (TR = 4200 ms; TE= 93 ms; field of view = 200 mm^2^; flip angle = 139°; bandwidth = 199 Hz/pixel; voxel size = 0.4 x 0.4 x 1.9 mm; 58 coronal slices acquired perpendicular to the long-axis of the hippocampus). High-resolution functional (T2*) images were acquired using a multiband gradient echo planar (EPI) imaging sequence (TR = 2010 ms; TE = 25 ms; field of view = 216 mm; image matrix = 144 x 152; flip angle = 79°; bandwidth = 1240 Hx/pixel; partial phase Fourier = 6/8; parallel imaging = GRAPPA acceleration factor 2 with 72 reference lines; multiband factor = 2; 52 oblique axial slices acquired parallel to the long-axis of the hippocampus slices; voxel size = 1.5 mm isotropic).

SPM8 (http://www.fil.ion.ucl.ac.uk/spm/) was used for image pre-processing. Functional EPI images were realigned to the first image and resliced. No slice timing correction was performed due to the acquisition of multiple simultaneous slices with the multiband sequence (capabilities to handle multiband timing do not exist in SPM8). Co-registration between the native-space ROIs defined in T2 space and the functional images was done with SPM’s Coregister: Estimate and Reslice procedure. This procedure uses a linear normalized mutual information cost-function between a reference (mean functional) image and source (T2) image to compute and apply a voxel-by-voxel affine transformation matrix. This transformation matrix was then applied to the subfield ROIs that had been defined in T2 space (see *ROI segmentation*) to bring them into register with the functional images. The T1 image was co-registered to the mean EPI. Then, nonlinear spatial normalization parameters were derived by segmenting the coregistered T1 image. Quality assurance included identifying suspect timepoints via custom code (https://github.com/memobc/memolab-fmri-qa) defined as time-points in excess of 0.5 mm frame displacement (based on (Power et al., 2012) or 1.5% global mean signal change (based on ARTRepair recommendations, (Mazaika et al., 2005)). Runs were excluded if the frame displacement exceeded the voxel size. As reported earlier, three participants were excluded for motion in excess of these thresholds; of the 24 subjects included in the analyses, 9 had runs excluded based on these thresholds (mean number of removed runs = 0.92, SD = 1.38; ranging from 0-4 runs).

### Pattern similarity analyses

PS analyses were conducted on beta maps generated from unsmoothed data in native subject space. Following the least squares separate procedure described by Mumford (2012), single trial models were generated to estimate the unique beta map for every trial in a run (N=60). Within each single trial model, the first regressor modeled the trial of interest with a stick function, the second regressor modeled all other trials in that run, six regressors were used to capture motion, and any additional spike regressors as identified by our QA scripts were used to capture additional residual variance. Voxel-wise patterns of hemodynamic activity were separately extracted for each ROI from the single trial beta images. To ensure robust ability to detect differences in PS, we required temporal signal-to-noise ratios (TSNR) in a region to be above 20 (approximately 2 standard deviations below the mean global TSNR of 50.4). This required the removal of entorhinal cortex and its subregions (mean TSNR ranged between 10-20), despite its compelling role in the representation of temporal context (Bellmund et al., 2019; Montchal et al., 2019)

Within each ROI, correlations (Pearson’s r) were computed between these trial-wise betas to yield a trial-by-trial correlation matrix that related each voxel’s signal on a trial to all other trials across all runs. We restricted comparisons to those trials for which participants made a correct “remember” response (during MRI scanning). Trials were sorted on the basis of encoding context (cognitive, temporal), however source memory for these contexts was not taken into consideration. Correlation values were z-transformed prior to statistical analysis. Statistical analyses tested for differences in correlations between trial pairs on the basis of encoding context (cognitive context: same vs. different encoding question; temporal context: same vs. different encoding list, or similar vs. different list half). To more accurately characterize both within- and across-subject error variance (Baayen et al., 2008; Clark, 1973; Dixon, 2008; Jaeger, 2008; Mumford & Poldrack, 2007; Singmann & Kellen, 2019), we implemented a mixed-modelling approach to evaluate statistical significance with the *lme4* packing in R (Bates et al., 2014); for a similar approach, see (Dimsdale-Zucker et al., 2018). We used an alpha of *p* < 0.05 and performed permutation testing for all of the pattern similarity results reported in the manuscript. Only between-run correlations were used to maximize the number of possible trial pairs without mixing within- and between-run correlations. Trial pairs of interest were extracted from these trial-by-trial correlation matrices.

All relevant code (https://github.com/hallez/tempcon_pub), a reproducible compute environment (https://doi.org/10.24433/CO.0129473.v1), and relevant data (https://osf.io/qfcjg/) are available online.

### ROI definition

Hippocampal subfields were defined following the procedure reported in (Dimsdale-Zucker et al., 2018; see Figure 2). In short, the ASHS automated segmentation procedure was used to delineate subfields in subject-native space (Yushkevich et al., 2010). We restricted our analyses to hippocampal body where discriminating subfields is most agreed upon. Medial temporal lobe cortical regions were manually-traced (see the Libby and Ranganath protocol in (Yushkevich et al., 2015)). In accordance with prior findings suggesting functional distinctions between anterior and posterior parahippocampal gyrus (Aminoff et al., 2007; Baldassano et al., 2013, 2016), we subdivided parahippocampal cortex one slice posterior to the wing of the ambient cistern (Frankó et al., 2014).

**Figure 2.**
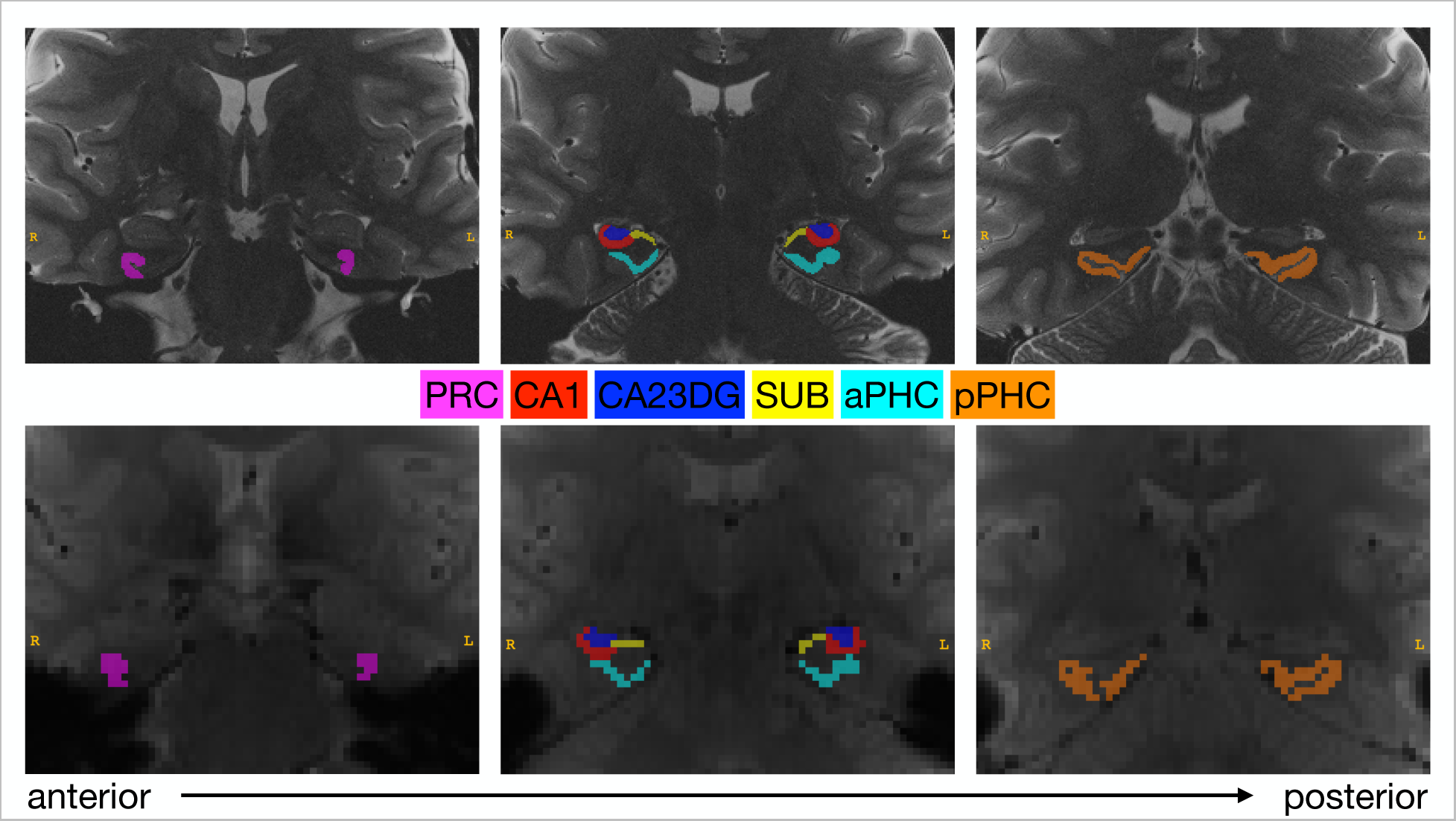
Top: Segmentations of hippocampal subfields (Cornu Ammonis [CA] 1, a combined CA2, CA3, and dentate gyrus [CA23DG] region, and subiculum [SUB]) as well as cortical medial temporal lobe regions (perirhinal cortex [PRC], anterior parahippocampal cortex [aPHC], and posterior parahippocampal cortex [pPHC]). Segmentations are depicted for a representative subject in the coronal plane of a T2 image. Slices move from anterior to posterior from left-to-right. Bottom: Segmentations following reslicing to functional (EPI) space displayed on a mean functional image from the same subject.

Because ROI definition and selection for comparisons was a priori, we performed independent analyses on each ROI. As Poldrack and Mumford (2009) suggest, taking this approach does not necessitate multiple comparison correction. Furthermore, we were primarily interested in characterizing the content of reactivated encoding contextual information within *each* subfield rather than making direct comparisons between the similarities or differences in what types of information each subfield primarily represented.

## Results

### Behavioral results

During MRI scanning participants performed a recognition memory test requiring judgments as to whether each item was recognized on the basis of recollection of specific item and source information from the study phase (see *Method*). Pattern similarity analyses were restricted to correctly remembered items. On average, this yielded 200 trials (SD = 51.9; minimum trials = 71; maximum trials = 282) from which we constructed all possible trial pairs and then compared activity patterns (see fMRI Results).

Correct remember judgments were the most common response (mean hit rate = 0.69, SD = 0.19), and, for these items, participants showed high accuracy at remembering the associated encoding task context (mean hit rate = 0.71, SD = 0.12). Memory for the exact (“same”) temporal context (list 1-8) was poor (mean hit rate = 0.18, SD = 0.03). Memory for task context and exact temporal context for items that were correctly remembered as studied were not significantly correlated with one another (r = -0.31, *p* = 0.16).

We reasoned that, even if participants were unable to recall the exact list identity, they might have memory for the “similar” temporal context associated with each item. Visual inspection of temporal source errors indicated that participants were more likely to incorrectly identify an item as coming from a list near in time to when it was actually studied as compared to lists further away (e.g., if an item was actually studied in List 3, participants would have been more likely to incorrectly say it had been in List 2 or 4 than lists further away in time; see Appendix A).

We therefore performed an exploratory analysis in which we re-scored each trial according to whether the participant could accurately determine whether it was presented in the “similar” (first half (lists 1-4) or the second half (lists 5-8)) versus “different” (across halves, i.e., list 1/list5, list 1/list 6, etc.) temporal context of the encoding phase. On this metric, memory for similar temporal context (mean accuracy for list half = 0.59, SD = 0.04) was reliably greater than chance of 0.5, t(22) = 8.74, *p* < 0.001. Thus, although participants did not have access to the precise list in which an item had been studied, they were able to retrieve information about the item’s temporal context at a coarse level.

**Appendix A.**
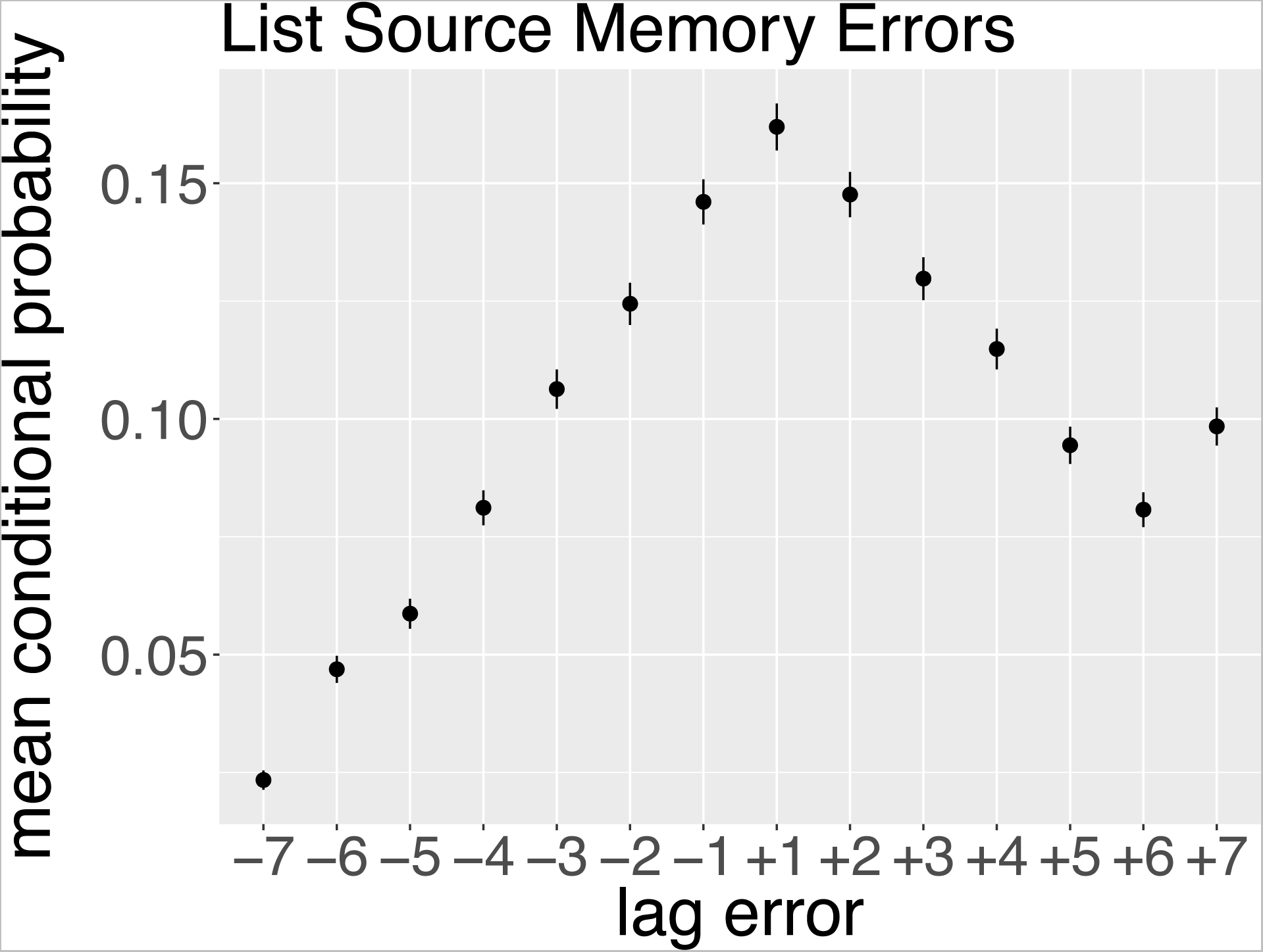
Temporal (list) source memory errors were more likely to occur at nearer than further lags. That is, items were more likely to be incorrectly attributed to a list that was closer in time (+/-1 or +/-2 lists) to the list in which it was actually located at encoding. We take this as evidence that although exact memory for which of the eight encoding lists an item was studied in was relatively poor, participants had access to the relative temporal context of an item at encoding.

### fMRI results

We next tested whether activity patterns in the hippocampal subfields (CA1, CA23DG, subiculum^1^) during memory retrieval carried information about the context in which the item was previously encountered. Specifically, we examined voxel PS during retrieval as a function of whether pairs of trials shared a temporal (same list vs. different list [*same* temporal context] and/or cognitive (i.e., same encoding task vs. different encoding task) context when the items were originally learned.

We first considered when items came from the same temporal encoding context (same vs. different list). No hippocampal subfield showed significant PS differences between retrieval of items from the same list as compared with items from different lists (all X^2^ < 0.5, all *p*s > 0.40; see Appendix C for statistics and Appendix B for visualization of PS values between lists).

**Appendix B.**
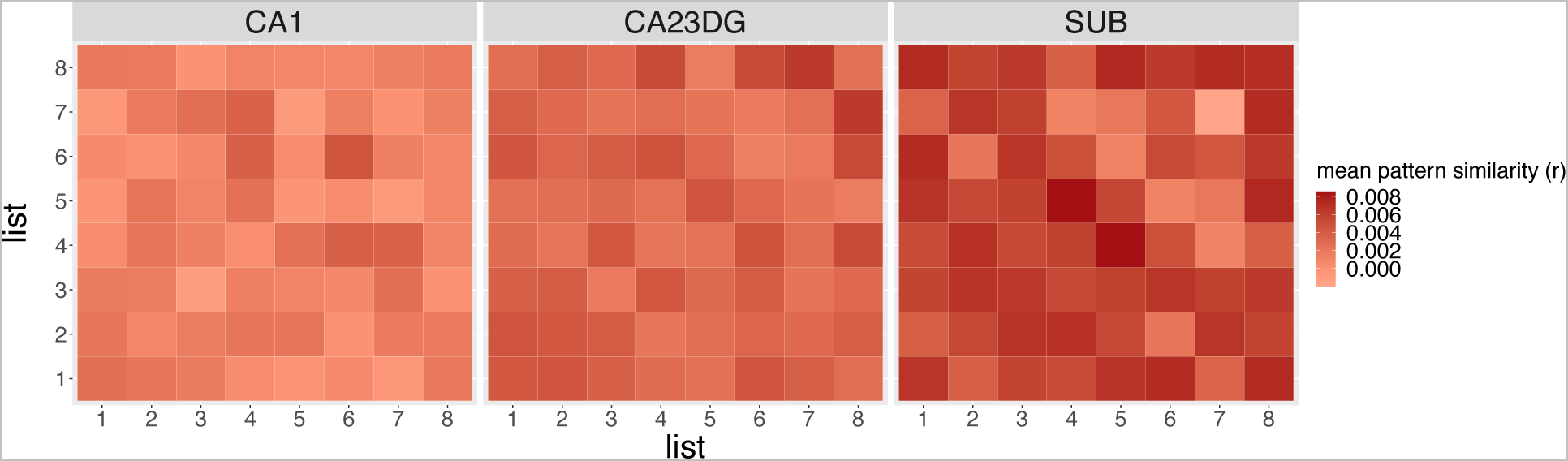
Pattern similarity split by list across subfields of interest. No subfield showed a reliable relationship in PS levels across list. We take this as evidence that list membership was not a salient feature that participants reactivated at the time of retrieval. This aligns with participants’ poor source memory for the list in which an item was studied during encoding.

We next considered whether hippocampal voxel patterns during recollection of studied items carried information about the cognitive context (encoding question) associated with the item during encoding. In CA23DG, we found that PS was higher during retrieval of items associated with the same cognitive context compared to items associated with different cognitive contexts (X^2^(1) = 4.63, p^perm1000^ = 0.031; Figure 3^2^). No other subfield showed significant effects of cognitive context (all X^2^ < 0.3, all *p*s > 0.50; see Appendix D) nor the combination of same versus different temporal and cognitive contexts (all X^2^ < 0.80, all *p*s > 0.30; see Appendix D) on PS^3^.

**Figure 3.**
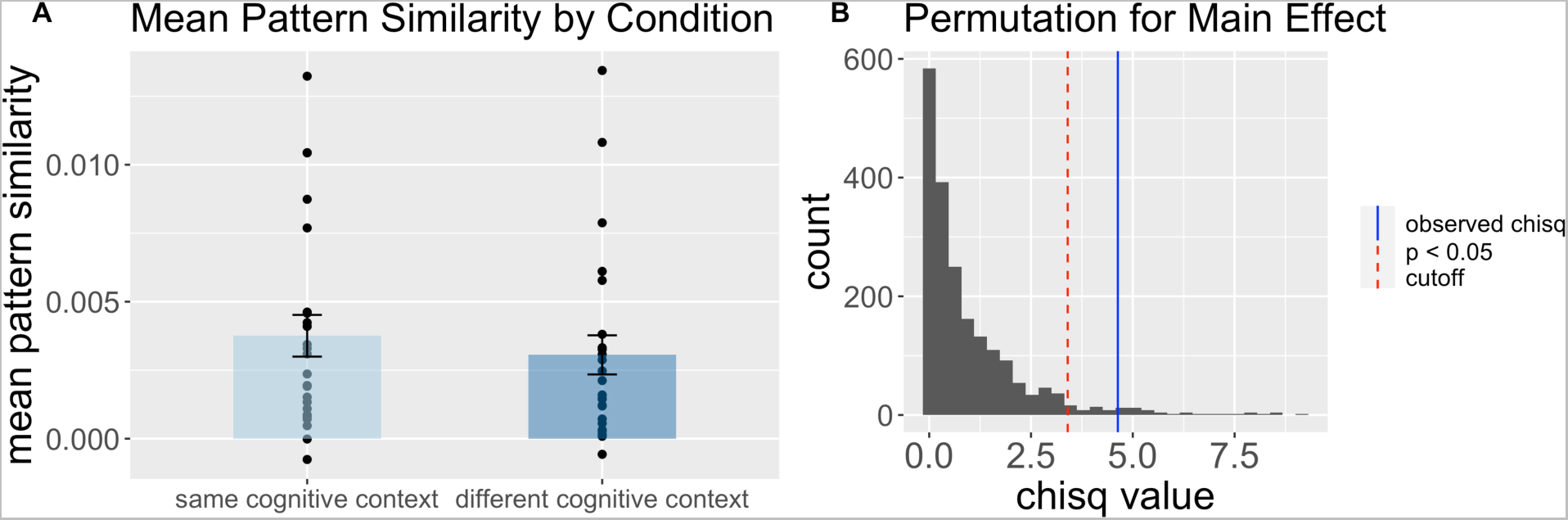
Pattern similarity values in CA23DG during memory retrieval carry information about cognitive encoding contexts. A. Mean pattern similarity scores, with scatter of individual subject observations for the combination of different encoding contexts. Mean PS values were greater for same as compared to different cognitive contexts. B. Permuted chi-square values to determine significance of cognitive context main effect. Observed chi-square value is depicted in blue and the significance threshold (*p* = 0.05) is plotted in a dashed red line.

We next performed a series of exploratory analyses to consider whether PS might depend on whether items shared a “similar” temporal context at encoding (same vs. different half of the encoding phase). The choice of list half was informed by the fact that participants were behaviorally able to discriminate coarse temporal context reliably greater than chance, however we present these analyses as exploratory because participants were not explicitly asked to make temporal source judgments at the granularity of list half. This analysis revealed no significant effects of similar temporal context alone (all X^2^ < 2.5, all *p*s > 0.10; see Appendix D).

We next considered whether hippocampal voxel patterns during recollection of studied items carried information about the cognitive context (encoding question) associated with the item during encoding when we defined temporal context as “similar”. Consistent with our previous analysis, in CA23DG, PS for pairs of items that were associated with the same cognitive context was higher than for pairs of items that were associated with different cognitive contexts (X^2^(1) = 4.68, p^perm1000^ = 0.031). This effect was qualified by a significant similar temporal by cognitive context interaction (X^2^(1) = 8.11, p^perm1000^ = 0.004; see Figure 4), such that the effect of cognitive context in CA23DG was larger for items that were in similar temporal contexts (same half) than for items that were in different temporal contexts (different half) particularly when these items shared the same cognitive context. No other subfield showed significant effects of cognitive context (all X^2^ < 0.30, all *p*s > 0.80; see Appendix D) nor the combination of similar versus different temporal and cognitive contexts (all X^2^ < 3, all *p*s > 0.080; see Appendix D) on PS.

**Figure 4.**
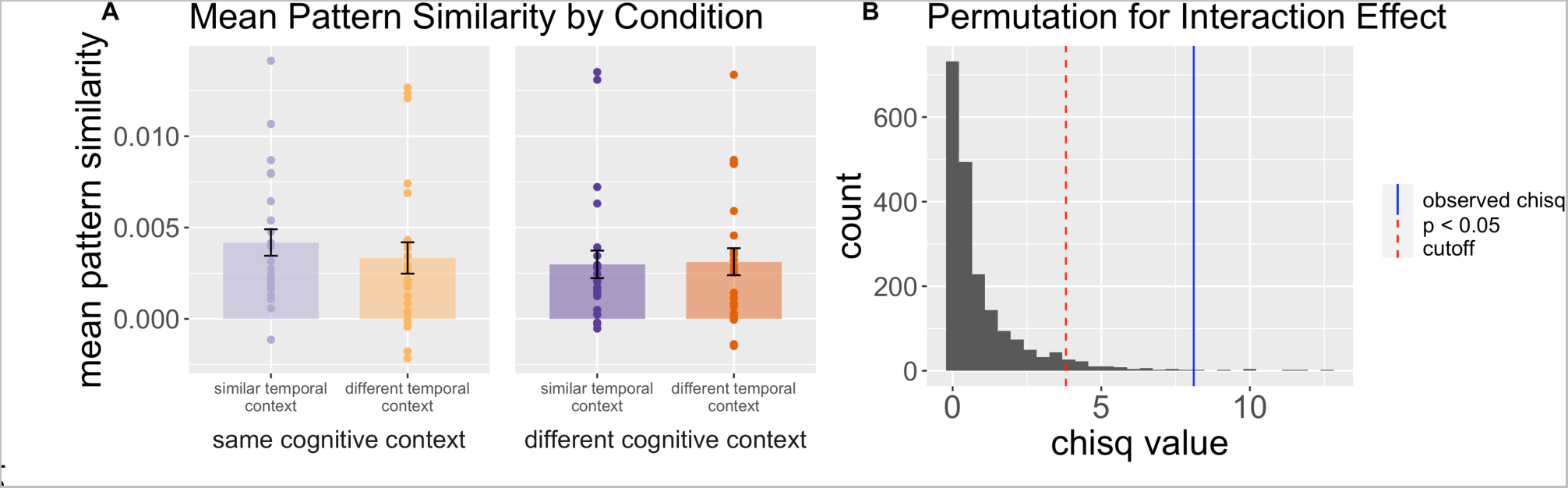
Pattern similarity values in CA23DG during memory retrieval carry information about similar temporal and cognitive encoding contexts. A. Mean pattern similarity scores, with scatter of individual subject observations for the combination of different encoding contexts. B. Permuted chi-square values to determine significance of cognitive-by-(similar)temporal interaction. Observed chi-square value is depicted in blue and the significance threshold (p = 0.05) is plotted in a dashed red line.

**Appendix C.** Pattern similarity findings in hippocampal subfields. Italics indicate significant modulation of pattern similarity values by cognitive, temporal, or interaction of cognitive and temporal encoding contexts.

|  | “Same” Temporal Context |  |  | “Similar” Temporal Context |  |  |
| --- | --- | --- | --- | --- | --- | --- |
|  | Cognitive | Temporal | Cognitive x Temporal | Cognitive | Temporal | Cognitive x Temporal |
| **CA1** | PS levels did not vary by cognitive context $X^2(1) = 0.08$, $p_{\text{perm1000}} = 0.773$ | PS levels did not vary by same/different temporal context $X^2(1) = 0.11$, $p_{\text{perm1000}} = 0.744$ | PS levels did not vary by the interaction between same temporal and cognitive contexts $X^2(1) = 0.03$, $p_{\text{perm1000}} = 0.871$ | PS levels did not vary by cognitive context ($X^2(1) = 0.08$, $p_{\text{perm1000}} = 0.776$) | PS levels did not vary by similar temporal context ($X^2(1) = 2.42$, $p_{\text{perm1000}} = 0.120$) | PS levels did not vary by the interaction between similar temporal and cognitive contexts ($X^2(1) = 0.84$, $p_{\text{perm1000}} = 0.360$) |
| **CA23DG** | *PS values were greater for same as compared to different cognitive contexts $X^2(1) = 4.63$, $p_{\text{perm1000}} = 0.031$* | PS levels did not vary by same/different temporal context $X^2(1) = 0.44$, $p_{\text{perm1000}} = 0.505$ | PS levels did not vary by the interaction between same temporal and cognitive contexts $X^2(1) = 0.71$, $p_{\text{perm1000}} = 0.400$ | *PS values were greater for same as compared to different cognitive contexts $X^2(1) = 4.68$, $p_{\text{perm1000}} = 0.031$* | PS levels did not vary by similar temporal context $X^2(1) = 0.08$, $p_{\text{perm1000}} = 0.783$ | *PS was greater for items that shared a similar temporal context as well as a cognitive context but was lower when these items did not share a cognitive context $X^2(1) = 8.11$, $p_{\text{perm1000}} = 0.004$* |
| **Subiculum** | PS levels did not vary by cognitive context $X^2(1) = 0.27$, $p_{\text{perm1000}} = 0.605$ | PS levels did not vary by same/different temporal context $X^2(1) = 0.13$, $p_{\text{perm1000}} = 0.721$ | PS levels did not vary by the interaction between same temporal and cognitive contexts $X^2(1) = 0.03$, $p_{\text{perm1000}} = 0.856$ | PS levels did not vary by cognitive context ($X^2(1) = 0.27$, $p_{\text{perm1000}} = 0.602$) | PS levels did not vary by similar temporal context ($X^2(1) = 0.11$, $p_{\text{perm1000}} = 0.745$) | PS was *marginally* higher for items that were studied relative to the same question but studied in different list halves as compared to all other conditions ($X^2(1) = 2.91$, $p_{\text{perm1000}} = 0.088$) |

**Appendix D.** Bayesian model results for pattern similarity findings in CA23DG. Here we report effect estimates for contrasts as estimated by the *brms* package in the R computing environment (Bürkner, 2017). Highest density intervals (abbreviated in the table as “posterior intervals” [PI]) were calculated for each contrast. Such intervals are roughly analogous to confidence intervals, and, thus, must not contain zero for the contrast to be considered significant (Turkkan & Pham-Gia, 1993). Italics are used to emphasize contrasts that were deemed significant based on this criteria.

|  | “Same” Temporal Context |  |  | “Similar” Temporal Context |  |  |
| --- | --- | --- | --- | --- | --- | --- |
|  | Cognitive | Temporal | Cognitive x Temporal | Cognitive | Temporal | Cognitive x Temporal |
| **CA23DG** | *PS values were greater for same as compared to different cognitive contexts; same > different cognitive context estimate = 0.0004, PI = (0.00005, 0.0008)* | PS levels did not vary by same/different temporal context; different > same list estimate = 0.000182, PI = (-0.0003, 0.0007) | PS levels did not vary by the interaction between same temporal and cognitive contexts; different list & same cognitive context > same list & different cognitive context estimate = -0.0005, PI = (-0.002, 0.00007) | *PS values were greater for same as compared to different cognitive contexts; same > different cognitive context estimate = 0.0004, PI = (0.00005, 0.0008)* | PS levels did not vary by similar temporal context; different half > same half estimate = 0.00005; PI = (-0.00003, 0.00004) | *PS was greater for items that shared a similar temporal context as well as a cognitive context but was lower when these items did not share a cognitive context. For items in different list halves, PS values were relatively similar regardless of same vs. different cognitive context; different list half & same cognitive context > same list half & different cognitive context estimate = -0.004, PI = (-0.002, -0.00003)* |

**Appendix E.**
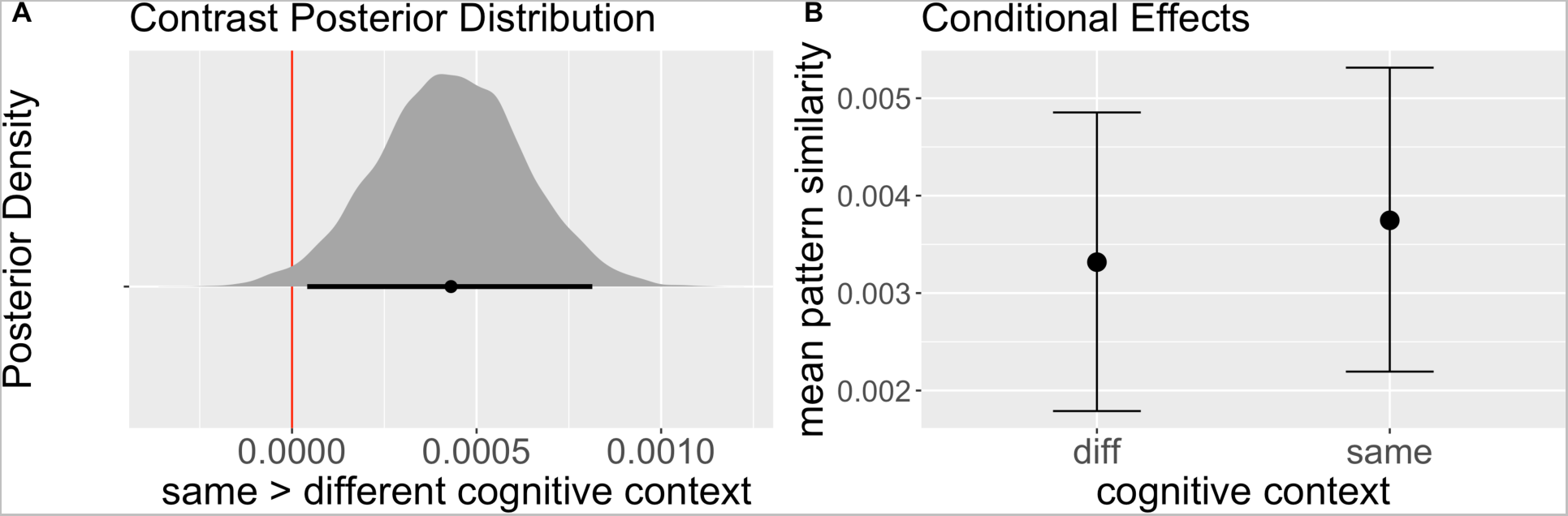
Bayesian model posterior distributions and conditional effects for the influence of cognitive context (for “same” temporal context items) on PS values in CA23DG. A. Posterior distributions for the same > different cognitive context contrast, representing differences in mean pattern similarity values between these conditions. Shaded distributions represent all posterior contrast samples, vertical red line represents zero, black point below shaded distribution represent effect estimates as determined by the model’s posterior predictive distribution for the linear predictor, and error bars around this point represent 95% credible intervals. Since the 95% CI does not overlap with zero, we can determine that cognitive context had a significant influence on PS values in CA23DG. B. Conditional effect estimates for the influence of same and different cognitive context on pattern similarity values, accounting for other effects present in the model.

**Appendix F.**
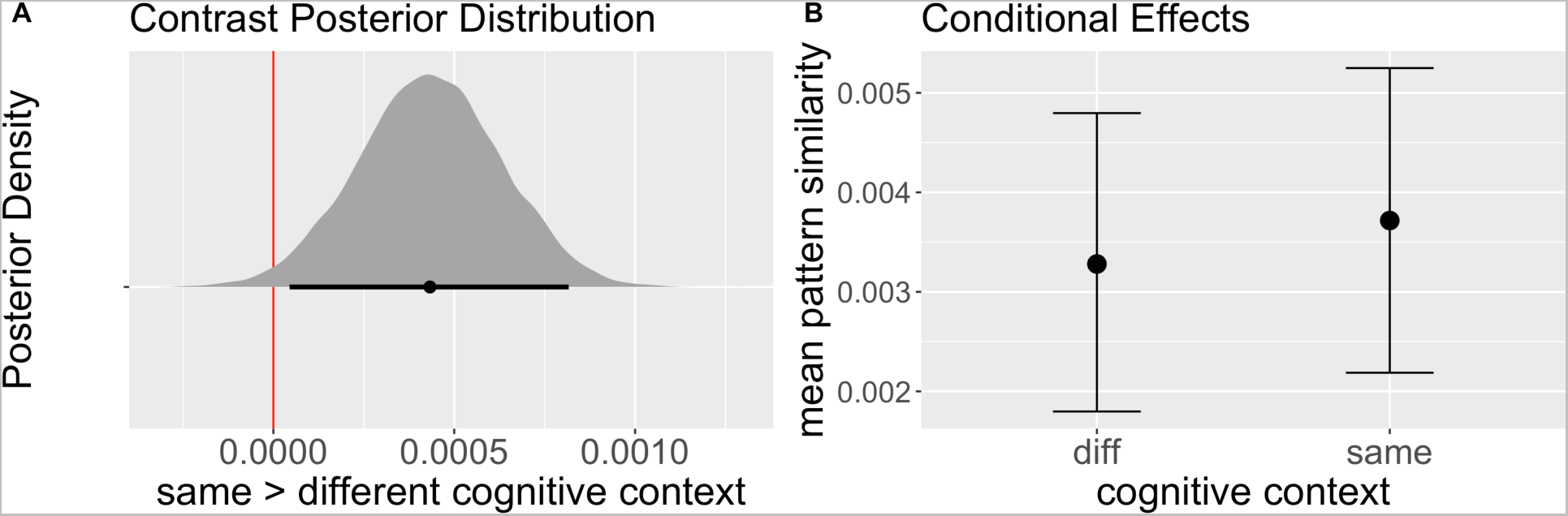
Bayesian model posterior distributions and conditional effects for the influence of cognitive context (for “similar” temporal context items) on PS values in CA23DG. A. Posterior distributions for the same > different cognitive context contrast, representing differences in mean pattern similarity values between these conditions. Shaded distributions represent all posterior contrast samples, vertical red line represents zero, black point below shaded distribution represent effect estimates as determined by the model’s posterior predictive distribution for the linear predictor, and error bars around this point represent 95% credible intervals. Since the 95% CI does not overlap with zero, we can determine that cognitive context had a significant influence on PS values in CA23DG. B. Conditional effect estimates for the influence of same and different cognitive context on pattern similarity values, accounting for other effects present in the model.

**Appendix G.**
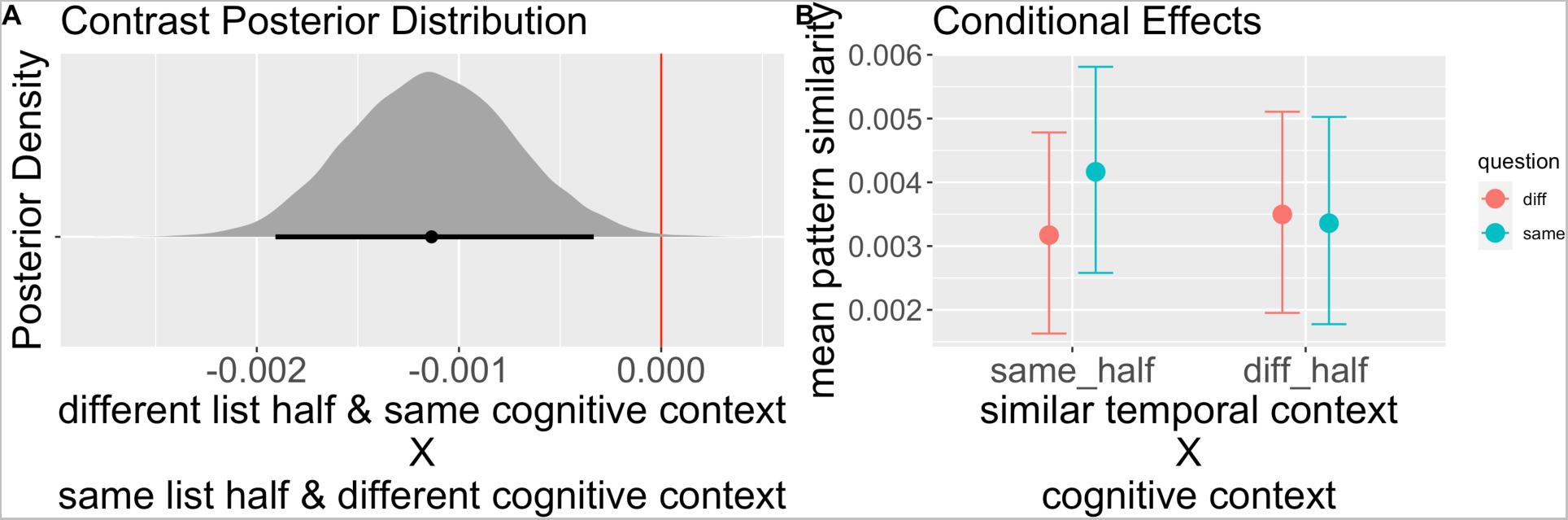
Bayesian model posterior distributions and conditional effects for the influence of the combined effect of “similar” temporal context and cognitive context on PS values in CA23DG. A. Posterior distributions for the [(different list half & same cognitive context) > (same list half & different cognitive context)] contrast, representing differences in mean pattern similarity values between these conditions. Shaded distributions represent all posterior contrast samples, vertical red line represents zero, black point below shaded distribution represent effect estimates as determined by the model’s posterior predictive distribution for the linear predictor, and error bars around this point represent 95% credible intervals. Since the 95% CI does not overlap with zero, we can determine that the combined influence of similar temporal context and cognitive context had a significant influence on PS values in CA23DG. Specifically, we see that PS values were greater for items affiliated both with the same cognitive context and the same list half (“similar” temporal context) than those affiliated with different cognitive contexts (comparing teal vs. red points in the left column of the graph). PS values were relatively equivalent for items affiliated with different list halves regardless of cognitive context (same/different). B. Conditional effect estimates for the influence of similar temporal context and cognitive context on pattern similarity values, accounting for other effects present in the model.

**Appendix H.**
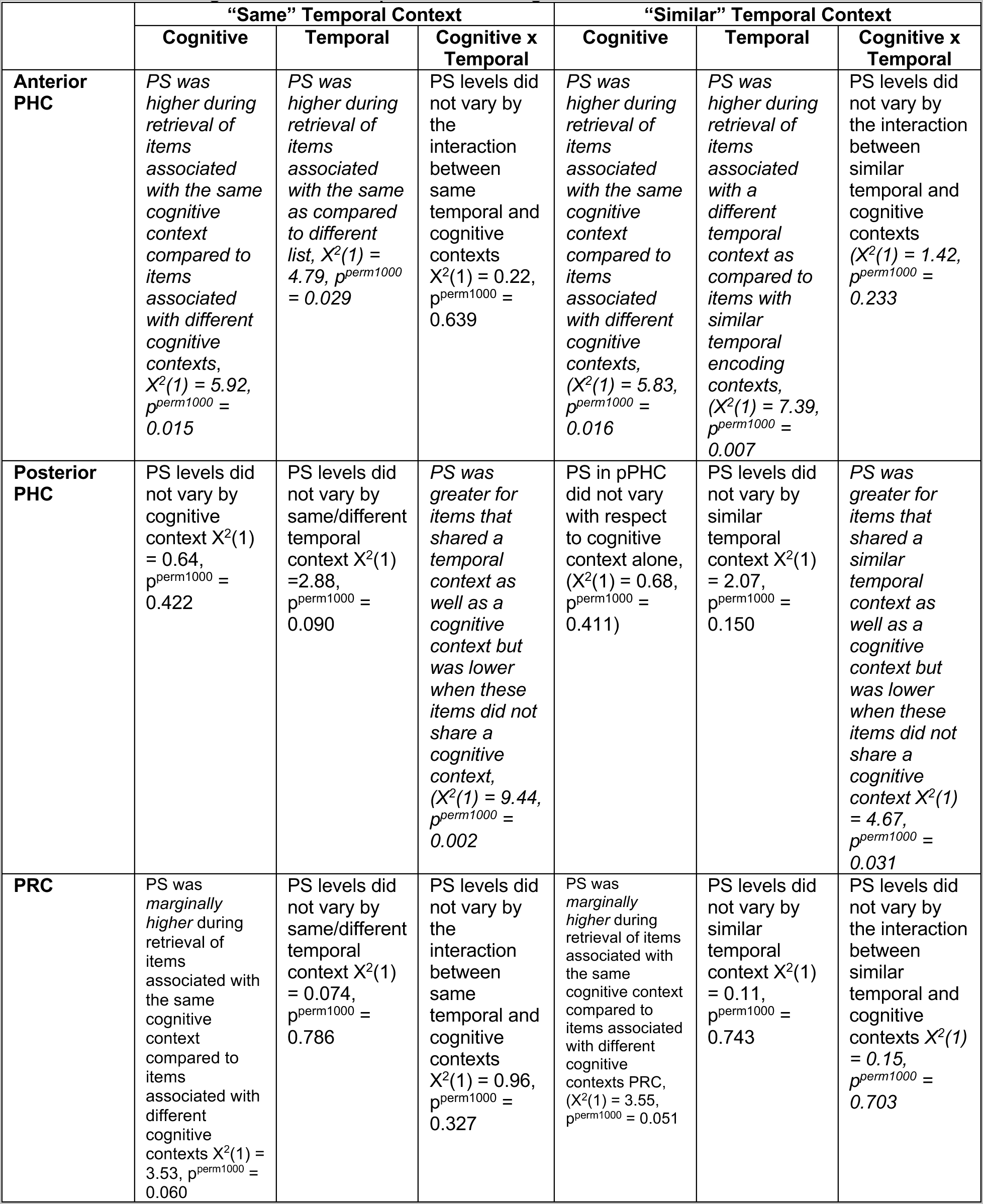
Pattern similarity findings in medial temporal lobe neocortical areas. Italics indicate significant modulation of pattern similarity values by cognitive, temporal, or interaction of cognitive and temporal encoding contexts.

|  | “Same” Temporal Context |  |  | “Similar” Temporal Context |  |  |
| --- | --- | --- | --- | --- | --- | --- |
|  | Cognitive | Temporal | Cognitive x Temporal | Cognitive | Temporal | Cognitive x Temporal |
| **Anterior PHC** | *PS was higher during retrieval of items associated with the same cognitive context compared to items associated with different cognitive contexts, $X^2(1) = 5.92$, $p^{\text{perm1000}} = 0.015$* | *PS was higher during retrieval of items associated with the same as compared to different list, $X^2(1) = 4.79$, $p^{\text{perm1000}} = 0.029$* | PS levels did not vary by the interaction between same temporal and cognitive contexts $X^2(1) = 0.22$, $p^{\text{perm1000}} = 0.639$ | *PS was higher during retrieval of items associated with the same cognitive context compared to items associated with different cognitive contexts, $(X^2(1) = 5.83$, $p^{\text{perm1000}} = 0.016$* | *PS was higher during retrieval of items associated with a different temporal context as compared to items with similar temporal encoding contexts, $(X^2(1) = 7.39$, $p^{\text{perm1000}} = 0.007$* | PS levels did not vary by the interaction between similar temporal and cognitive contexts $(X^2(1) = 1.42$, $p^{\text{perm1000}} = 0.233$ |
| **Posterior PHC** | PS levels did not vary by cognitive context $X^2(1) = 0.64$, $p^{\text{perm1000}} = 0.422$ | PS levels did not vary by same/different temporal context $X^2(1) = 2.88$, $p^{\text{perm1000}} = 0.090$ | *PS was greater for items that shared a temporal context as well as a cognitive context but was lower when these items did not share a cognitive context, $(X^2(1) = 9.44$, $p^{\text{perm1000}} = 0.002$* | PS in pPHC did not vary with respect to cognitive context alone, $(X^2(1) = 0.68$, $p^{\text{perm1000}} = 0.411$) | PS levels did not vary by similar temporal context $X^2(1) = 2.07$, $p^{\text{perm1000}} = 0.150$ | *PS was greater for items that shared a similar temporal context as well as a cognitive context but was lower when these items did not share a cognitive context $X^2(1) = 4.67$, $p^{\text{perm1000}} = 0.031$* |
| **PRC** | PS was *marginally higher* during retrieval of items associated with the same cognitive context compared to items associated with different cognitive contexts $X^2(1) = 3.53$, $p^{\text{perm1000}} = 0.060$ | PS levels did not vary by same/different temporal context $X^2(1) = 0.074$, $p^{\text{perm1000}} = 0.786$ | PS levels did not vary by the interaction between same temporal and cognitive contexts $X^2(1) = 0.96$, $p^{\text{perm1000}} = 0.327$ | PS was *marginally higher* during retrieval of items associated with the same cognitive context compared to items associated with different cognitive contexts PRC, $(X^2(1) = 3.55$, $p^{\text{perm1000}} = 0.051$ | PS levels did not vary by similar temporal context $X^2(1) = 0.11$, $p^{\text{perm1000}} = 0.743$ | PS levels did not vary by the interaction between similar temporal and cognitive contexts $X^2(1) = 0.15$, $p^{\text{perm1000}} = 0.703$ |

## Discussion

The goal of the present study was to test whether hippocampal subfields represent information about experimentally-manipulated cognitive and temporal contexts (i.e., encoding task and list identity) during recollection of studied items. Results showed that cognitive context significantly influenced activity patterns in CA23DG. We saw no evidence for a precise representation of temporal (list) context, although exploratory analyses revealed that when using a coarse definition of temporal context (first vs second half of study phase), that CA23DG carried information about the *conjunction* of cognitive and temporal context. This suggests that patterns of activity within CA23DG may support successful memory retrieval by enabling recollection of task-relevant cognitive states or goals.

### Cognitive Context Representation in CA23DG

The hippocampus is a prime example of how form can influence function. Computational models (Marr, 1971; for a non-Hebbian instantiation, see Y. Zheng et al., 2021) propose that the combination of sparse coding in DG and the dense recurrent collaterals that make up the majority of the inputs to CA3 (Amaral & Witter, 1989) enable the hippocampus to retrieve or recover a memory trace given a noisy or incomplete retrieval cue (i.e., “pattern completion” (Marr, 1971; O’Reilly & McClelland, 1994; Yassa & Stark, 2011). A second factor that is key to understanding hippocampal function is that CA3 and DG are critical sites for binding of information about items and contexts (Knierim et al., 2006; Y. Zheng et al., 2021). Considerable evidence suggests that lateral and medial entorhinal cortex receive distinct cortical inputs carrying information about items and contexts, respectively, and this information is subsequently integrated via convergent inputs in DG and CA3 (Amaral & Witter, 1989; Burwell, 2000; Eichenbaum et al., 2007; Libby et al., 2012; Rolls & Kesner, 2006). Based on these characteristics, we would expect that the DG-CA3 circuit should play a crucial role in recollecting contextual information about previously encountered items.

Using a paradigm adapted from Diana and colleagues (2012, 2013), in which different objects were encoded with unique cognitive context questions, we demonstrated that activity patterns in CA23DG carried information about an item’s cognitive context during retrieval. Pattern similarity values were greater for items that had been affiliated with the same, as compared to different, cognitive tasks at encoding. This result is striking, given that participants were only shown items from the study phase, and they were not instructed to explicitly recall information about the study tasks. Our results therefore accord with the idea that CA23DG plays an important role in linking item and context information, such that encountering a familiar item can trigger the recovery of contextual information via pattern completion (see also Grande et al., 2019).

Our findings are relevant to the idea that, during conscious recollection, the hippocampus reinstates representations of discrete cognitive contexts that encompass one’s current attentional priorities, task set, and/or goals (Aly & Turk-Browne, 2016a, 2016b; Davachi, 2006; Davachi et al., 2003; Diana et al., 2012; Eichenbaum et al., 2007; J. D. Johnson et al., 2009; Libby et al., 2018; Ranganath et al., 2004; Ritchey et al., 2015). For instance, Aly and colleagues (Aly & Turk-Browne, 2016a, 2016b) found that voxel patterns in hippocampal subfields carried information about attentional orientation during a memory encoding task (“attend to artistic features” vs. “attend to spatial layout”). Converging evidence in rodents also suggests that task-relevant contextual features were the most salient factor in describing hippocampal ensemble activity similarity (conceptually analogous to the pattern similarity metrics used in human fMRI) (McKenzie et al., 2014). Our results build on this idea by highlighting CA23DG as a site where binding of cognitive contextual information occurs in the service of episodic memory retrieval.

An intriguing possibility that could be addressed in future work is whether representational patterns would change if cognitive contexts were designed to systematically relate to features of the items. In the present design, item/question pairing was uniquely randomized for each participant and thus this question cannot be explored here. However, prior work has suggested that the hippocampus may play a particular role in representing an item’s context (e.g., the type of associated judgment) whereas when context is posed as a feature of the item (e.g., its color) that medial temporal cortical regions such as perirhinal cortex are engaged (Staresina & Davachi, 2008; although see Diana, Yonelinas, & Ranganath, 2008 for some suggestions that perirhinal involvement in item/context unitization may be limited to familiarity). Thus, if cognitive context questions were selected to represent *features* of the items, it seems that there would be competing hypotheses about whether hippocampus would still treat this information as an item’s context or whether medial temporal lobe cortical regions would additionally be recruited.

### Contributions of temporal context

According to Tulving’s definition (Tulving, 1972, 1983, 1984), episodic memories are organized by temporal context. Temporal context has been formally operationalized in terms of random fluctuations in cognitive states over time (Estes, 1955), and a time-weighted average of recently processed items and experiences (Howard & Kahana, 2002; Norman et al., 2008; Sederberg et al., 2008). These models would predict that activity patterns during memory retrieval should reflect information about the temporal context associated with each study item (e.g., Deuker et al., 2016; Jenkins & Ranganath, 2016; Manning et al., 2011; Nielson et al., 2015).

In the present study, when asked to choose in which of eight lists an item had been studied, participants were essentially at chance. This is not surprising because models suggest that temporal context is relative, not absolute (Kahana, 1996; Manning et al., 2011). In other words, people can use recovered context to gauge the relative recency of two previously encountered items but this might not be sufficient for precise memory for the temporal position of an item (DuBrow & Davachi, 2013, 2014; Ezzyat & Davachi, 2014; Hintzman, 2001, 2004, 2005; Jenkins & Ranganath, 2010, 2016; Lewandowsky & Murdock, 1989; Montchal et al., 2019) without the use of additional heuristics and reconstructive strategies (Friedman, 1993).

If the hippocampus carries information about the relative temporal context of past events, we might expect that hippocampal activity patterns would be more similar during recollection of items that were learned in close temporal proximity than for items that were studied far apart in time. Consistent with this prediction, fMRI studies have shown that recollection of items encountered either in virtual reality (Copara et al., 2014; Deuker et al., 2016; Dimsdale-Zucker et al., 2018) or in the real-world (Nielson et al., 2015), yield activity patterns within the hippocampus that show graded similarity on the basis of both temporal and spatial contextual features.

Here, we found no evidence to suggest that activity patterns in any subfield carried information about list context. There are a number of potential reasons for this null result, but we suspect that the most important factor is that, unlike previous studies that reported temporal context representations in the hippocampus, we manipulated cognitive contexts independently of temporal information.

Our task was designed such that participants changed cognitive contexts often within a list. Prior work has shown that when ongoing temporal contexts are interrupted by changes in item-level cognitive contexts that both behavioral (Polyn et al., 2009) and neural (Polyn et al., 2012) responses are shaped by both these local and global contextual features. Thus, by varying the encoding question within each study list, participants might have segmented each list into micro-contexts (Clewett & Davachi, 2017; DuBrow et al., 2017; Zacks et al., 2001; Zacks & Swallow, 2007) thereby disrupting associations based on temporal contiguity. A related possibility is that, during encoding, participants might have prioritized cognitive context over temporal contiguity, because the encoding questions were more salient and relevant.

Although we did not find evidence of precise temporal context representation in the hippocampus, an exploratory analysis did reveal evidence to suggest conjunctive representation of temporal context—“similar” or “coarse” temporal context—and cognitive context in CA23DG. In this analysis, temporal context was operationalized on the basis of which half of the study phase that an item had been previously encountered (first half: lists 1-4; second half: lists 5-8). Results revealed that activity patterns in CA23DG were shaped by the *conjunction* of coarse temporal context and cognitive context, which aligns with the binding role CA3 is thought to play (Burwell, 2000; Eichenbaum et al., 2007; Rolls & Kesner, 2006).

### Shared versus distinct contextual representations in the hippocampus

Our results conflict with a prior study, in which, we found that CA23DG activity patterns during memory retrieval were less similar across pairs of items that were associated with the same episodic context (i.e., objects seen within the same movie) than across pairs of items that were associated with different spatial or episodic contexts (Dimsdale-Zucker et al., 2018). The contrast between the present results and those of Dimsdale-Zucker et al. (2018) is relevant to many standard resolution fMRI studies showing that that hippocampal activity patterns are sometimes more similar across events that have a shared context (Horner et al., 2015; Libby et al., 2018; Milivojevic et al., 2015; Pidgeon & Morcom, 2016; Schlichting et al., 2015), and sometimes less similar between overlapping events (Chanales et al., 2017; Favila et al., 2016; Kim et al., 2017; Schlichting et al., 2015; L. Zheng et al., 2021). Computational work by Ritvo (2019) suggests that these different results may be due to competition between items that are to be learned. If there is relatively low competition, then CA23DG might assign similar representations to overlapping events, whereas, if competition is high, the representations can be orthogonalized. We speculate that dynamic might be at play in our studies. In Dimsdale-Zucker et al. (2018), multiple sequences of items were encoded in the same two virtual reality contexts, potentially creating a significant degree of interference across items, but, in the present study, the use of multiple encoding tasks probably reduced contextual overlap across items, thereby reducing competition. This explanation can be directly tested in a future high-resolution imaging study.

### General Conclusions

Our results show that activity patterns in the CA23DG region of the human hippocampus carry information about cognitive contexts. This finding can help explain how we represent continuously unfolding episodes in a changing world. Outside of the laboratory, we are constantly multitasking between competing goals and responsibilities. The hippocampus, and, specifically CA23DG, may allow us to differentiate between experiences that are associated with different tasks.

## Gender, Race, and Ethnicity Bias Statement

Recent work in several fields of science has identified a bias in citation practices such that papers from women and other minority scholars are under-cited relative to the number of such papers in the field (Bertolero et al., 2020; Caplar et al., 2017; Chatterjee & Werner, 2021; Dion et al., 2018; Dworkin et al., 2020; Fulvio et al., 2021; Maliniak et al., 2013; Mitchell et al., 2013; Wang et al., 2021). Here we sought to proactively consider choosing references that reflect the diversity of the field in thought, form of contribution, gender, race, ethnicity, and other factors. First, we obtained the predicted gender of the first and last author of each reference by using databases that store the probability of a first name being carried by a woman (Dworkin et al., 2020; Zhou, 2020/2022; Zhou et al., 2020). By this measure (and excluding self-citations to the first and last authors of our current paper), our references contain 7.65% woman(first)/woman(last), 14.53% man/woman, 26.29% woman/man, and 51.53% man/man. This method is limited in that a) names, pronouns, and social media profiles used to construct the databases may not, in every case, be indicative of gender identity and b) it cannot account for intersex, non-binary, or transgender people. Second, we obtained predicted racial/ethnic category of the first and last author of each reference by databases that store the probability of a first and last name being carried by an author of color (Ambekar et al., 2009; Sood & Laohaprapanon, 2018). By this measure (and excluding self-citations), our references contain 3.36% author of color (first)/author of color(last), 10.63% white author/author of color, 15.17% author of color/white author, and 70.84% white author/white author. This method is limited in that a) names and Florida Voter Data to make the predictions may not be indicative of racial/ethnic identity, and b) it cannot account for Indigenous and mixed-race authors, or those who may face differential biases due to the ambiguous racialization or ethnicization of their names. We look forward to future work that could help us to better understand how to support equitable practices in science.

## Footnotes

1 For medial temporal neocortical regions, see reported results in Appendix H.

2 For ease of visualization of the effects, for pattern similarity analyses we have chosen to plot means with individual subject points to illustrate the observed data. This does not exactly recapitulate the way the statistical comparisons were computed since these were performed as mixed models. That is, these figures show the raw pattern similarity values for these experimental conditions but do not account for the other effects present in the models that were used to determine significance. To better visualize the observed statistical effects, we have plotted the permutation distributions that were used to determine model significance.

3 At the suggestion of an anonymous reviewer, we repeated all of our analyses for the key effects we saw in CA23DG using a Bayesian modelling framework. We fit all models using Hamiltonian Monte Carlo No-U-Turn sampling as implemented by the brms package in the R computing environment (Bürkner, 2017). For all models, we fit 4 chains of 10,000 sampling iterations (5,000 warmup) each for a total of 20,000 post-warmup samples. This sampling iteration size was used to avoid cases where the tail effective sample size was low (as indicated by Stan warning messages). Extraction and transformation of posterior draws after models were fit was done using the *tidybayes* package and the *tidyverse* collection of packages in R (Kay, 2022; Wickham et al., 2019). This approach was informed by best practices as advised by Paul Alexander Bloom (personal communication). Results from these Bayesian analyses replicated the results we report in the main analyses in the paper that were implemented via mixed models in *lmer*. However, for full transparency, we have included the Bayesian effect estimates for all contrasts as well as the Highest Density Interval (which can be interpreted similarly to a confidence interval, Turkkan & Pham-Gia, 1993) in Appendix D. Furthermore, we include graphical depictions of the model posterior distribution and conditional effect estimates for any contrast that was deemed significant (see Appendices E-G).

## Acknowledgements.

The authors wish to thank Paul Alexander Bloom for his help in fitting and interpreting the Bayesian models.

